# Cyclooxgenase-2 is induced by SARS-CoV-2 infection but does not affect viral entry or replication

**DOI:** 10.1101/2020.09.24.312769

**Authors:** Jennifer S. Chen, Mia Madel Alfajaro, Jin Wei, Ryan D. Chow, Renata B. Filler, Stephanie C. Eisenbarth, Craig B. Wilen

## Abstract

Identifying drugs that regulate severe acute respiratory syndrome coronavirus 2 (SARS-CoV-2) infection and its symptoms has been a pressing area of investigation during the coronavirus disease 2019 (COVID-19) pandemic. Nonsteroidal anti-inflammatory drugs (NSAIDs), which are frequently used for the relief of pain and inflammation, could modulate both SARS-CoV-2 infection and the host response to the virus. NSAIDs inhibit the enzymes cyclooxygenase-1 (COX-1) and cyclooxygenase-2 (COX-2), which mediate the production of prostaglandins (PGs). PGE_2_, one of the most abundant PGs, has diverse biological roles in homeostasis and inflammatory responses. Previous studies have shown that NSAID treatment or inhibition of PGE_2_ receptor signaling leads to upregulation of angiotensin-converting enzyme 2 (ACE2), the cell entry receptor for SARS-CoV-2, thus raising concerns that NSAIDs could increase susceptibility to infection. COX/PGE_2_ signaling has also been shown to regulate the replication of many viruses, but it is not yet known whether it plays a role in SARS-CoV-2 replication. The purpose of this study was to dissect the effect of NSAIDs on COVID-19 in terms of SARS-CoV-2 entry and replication. We found that SARS-CoV-2 infection induced COX-2 upregulation in diverse human cell culture and mouse systems. However, suppression of COX-2/PGE_2_ signaling by two commonly used NSAIDs, ibuprofen and meloxicam, had no effect on *ACE2* expression, viral entry, or viral replication. Our findings suggest that COX-2 signaling driven by SARS-CoV-2 may instead play a role in regulating the lung inflammation and injury observed in COVID-19 patients.

**Importance:** Public health officials have raised concerns about the use of nonsteroidal anti-inflammatory drugs (NSAIDs) for treating symptoms of coronavirus disease 2019 (COVID-19), which is caused by severe acute respiratory syndrome coronavirus 2 (SARS-CoV-2). NSAIDs function by inhibiting the enzymes cyclooxygenase-1 (COX-1) and cyclooxygenase-2 (COX-2). These enzymes are critical for the generation of prostaglandins, lipid molecules with diverse roles in maintaining homeostasis as well as regulating the inflammatory response. While COX-1/COX-2 signaling pathways have been shown to affect the replication of many viruses, their effect on SARS-CoV-2 infection remains unknown. We found that SARS-CoV-2 infection induced COX-2 expression in both human cell culture systems and mouse models. However, inhibition of COX-2 activity with NSAIDs did not affect SARS-CoV-2 entry or replication. Our findings suggest that COX-2 signaling may instead regulate the lung inflammation observed in COVID-19 patients, which is an important area for future studies.

## Introduction

During the ongoing coronavirus disease 2019 (COVID-19) pandemic, a common concern has been whether widely used anti-inflammatory medications affect the risk of infection by severe acute respiratory syndrome coronavirus 2 (SARS-CoV-2), the causative agent of COVID-19, or disease severity. Used ubiquitously for the relief of pain and inflammation, nonsteroidal anti-inflammatory drugs (NSAIDs) have been one such target of concern, with the health minister of France and the medical director of the National Health Service of England recommending the use of acetaminophen over NSAIDs for treating COVID-19 symptoms (1, 2).

NSAIDs function by inhibiting the cyclooxygenase (COX) isoforms COX-1 and COX-2. COX-1 is constitutively expressed in most cells, while COX-2 expression is induced by inflammatory stimuli (3). COX-1 and COX-2 metabolize arachidonic acid into prostaglandin H_2_, which can then be converted to several different bioactive prostaglandins (PGs) (3). Prostaglandin E_2_ (PGE_2_) is one of the most abundant PGs in the body and signals through four receptors (EP1, EP2, EP3, and EP4) to perform diverse roles, such as regulating immune responses and gastrointestinal barrier integrity (3). Several potential hypotheses have linked NSAID use to COVID-19 pathogenesis. First, it has been suggested that NSAID use may upregulate angiotensin-converting enzyme 2 (ACE2), the cell entry receptor for SARS-CoV-2, and increase the risk of infection (4, 5). Second, given their anti-inflammatory properties, NSAIDs may impair the immune response to SARS-CoV-2 and delay disease resolution (1). Third, NSAIDs may also directly affect SARS-CoV-2 replication, as COX/PGE_2_ signaling has been shown to regulate replication of other viruses including mouse coronavirus (6). Therefore, given the widespread use of NSAIDs, evaluation of the interaction between NSAIDs and SARS-CoV-2 entry and replication is warranted.

NSAIDs may modulate multiple stages of the SARS-CoV-2 life cycle. As described above, one potential mechanism is that NSAIDs could lead to ACE2 upregulation and thus increase the susceptibility to SARS-CoV-2. Ibuprofen treatment of diabetic rats was found to increase ACE2 expression in the heart (7), though it was not studied whether the same occurs in non-diabetic rats. In addition, inhibition of the PGE_2_ receptor EP4 in human and mouse intestinal organoids increases ACE2 expression (8), suggesting that NSAID inhibition of COX/PGE_2_ signaling could similarly lead to ACE2 upregulation. NSAIDs could also affect a later stage of the SARS-CoV-2 life cycle. For porcine sapovirus, feline calicivirus, murine norovirus, and mouse coronavirus, COX inhibition impaired viral replication (6, 9, 10). COX inhibition was found to impair mouse coronavirus infection at a post-binding step early in the replication cycle, potentially entry or initial genome replication (6). Furthermore, SARS-CoV, the closest relative of SARS-CoV-2 among human coronaviruses and cause of the 2002-2003 epidemic (11), stimulates COX-2 expression via its spike protein and nucleocapsid protein (12, 13), indicating the potential relevance of this pathway for SARS-CoV-2.

Here, we assessed the relevance of COX-2/PGE_2_ signaling and inhibition by NSAIDs for SARS-CoV-2 infection. We found that SARS-CoV-2 infection induces COX-2 expression in human cells and mice. However, suppression of COX-2/PGE_2_ signaling by two commonly used NSAIDs, ibuprofen and meloxicam, had no effect on *ACE2* expression, viral entry, or viral replication. Together, this suggests that NSAID use in humans is unlikely to have adverse effects on SARS-CoV-2 transmission or pathogenesis.

## Results

To determine whether the COX-2/PGE_2_ pathway is relevant for SARS-CoV-2 infection, we evaluated induction of *PTGS2* (encoding COX-2) in human cells and mice. We found that SARS-CoV-2 infection of human lung cancer cell line Calu-3 led to significant upregulation of *PTGS2* (Fig. 1A). This is consistent with RNA sequencing (RNA-seq) datasets of SARS-CoV-2-infected Calu-3 cells and ACE2-overexpressing A549 cells, another lung cancer cell line (Fig. 1B-C) (14). However, infection of human liver cancer cell line Huh7.5 did not lead to significant *PTGS2* induction, demonstrating cell type specificity of *PTGS2* induction by SARS-CoV-2 (Fig. 1D).

**Figure 1.**
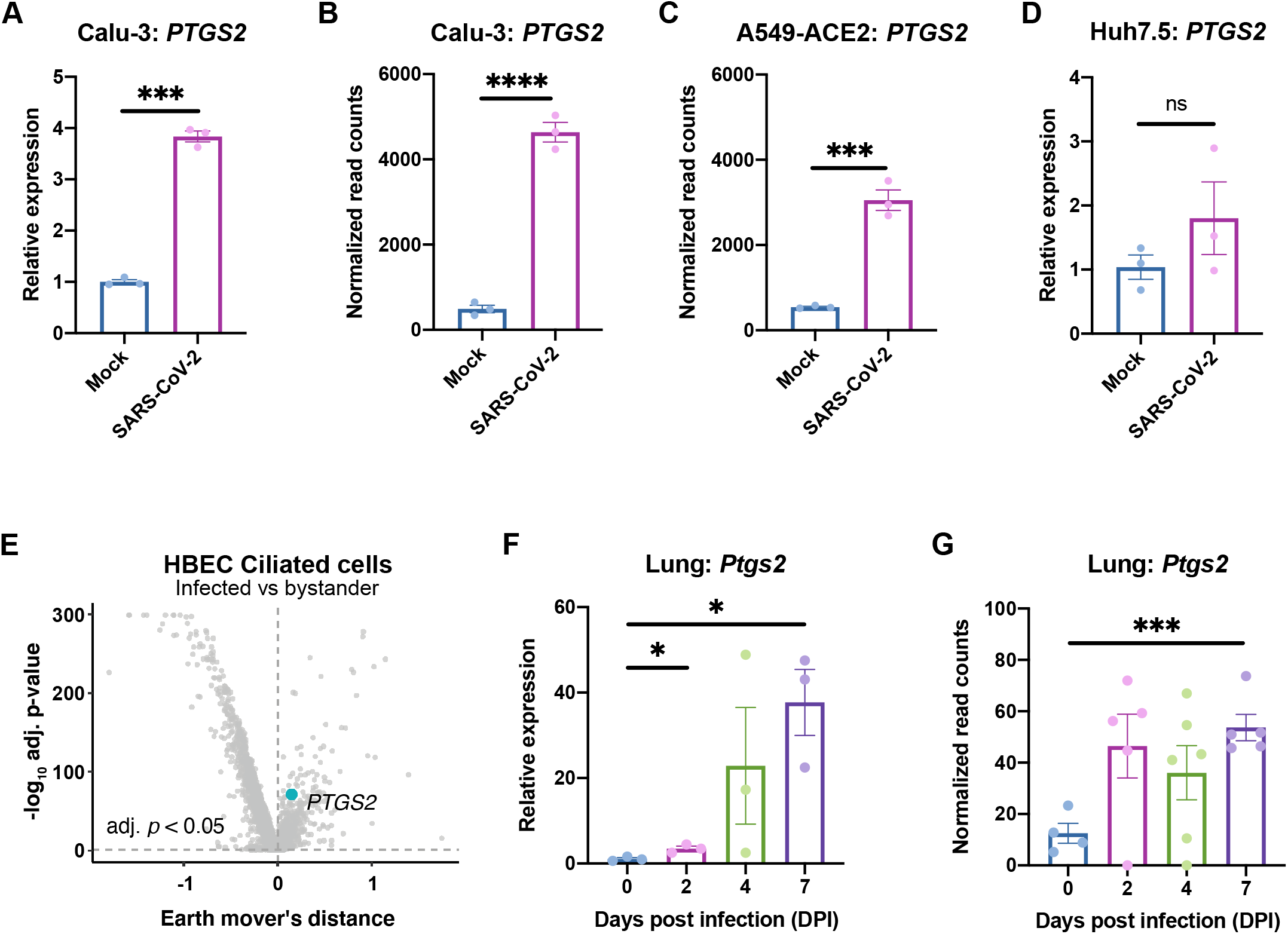
SARS-CoV-2 infection induces *PTGS2* expression in human and mouse systems. (A) Calu-3 cells were infected with SARS-CoV-2 at a multiplicity of infection (MOI) of 0.05. *PTGS2* expression was measured at 2 days post infection (dpi), normalized to *ACTB*. (B-C) *PTGS2* expression in Calu-3 (B) and ACE2-overexpressing A549 (A549-ACE2) (C) cells following SARS-CoV-2 infection. Data are from GSE147507 (Blanco-Melo et al., 2020). (D) Huh7.5 cells were infected with SARS-CoV-2 at a MOI of 0.05. *PTGS2* expression was measured at 2 dpi, normalized to *ACTB*. (E) Human bronchial epithelial cells (HBECs) were cultured at an air-liquid interface and then infected at the apical surface with 10^4^ plaque-forming units (PFU) of SARS-CoV-2. Cells were collected at 1, 2, and 3 dpi for single-cell RNA sequencing (scRNA-seq) (16). Volcano plot of differentially expressed genes in infected versus bystander ciliated cells pooled from all time points. *PTGS2* is highlighted. (F) K18-hACE2 mice were infected intranasally with 1.2 × 10^6^ PFU of SARS-CoV-2. *Ptgs2* expression in the lung was measured at 0, 2, 4, and 7 dpi. (G) *Ptgs2* expression in the lung of K18-hACE2 mice following intranasal SARS-CoV-2 infection. Data are from GSE154104 (Winkler et al., 2020). All data points in this figure are presented as mean ± SEM. Data were analyzed by Welch’s two-tailed, unpaired *t*-test (A, D, F); Student’s two-tailed, unpaired *t*-test (B, C, G); and two-sided Mann-Whitney U test with continuity and Benjamini-Hochberg correction (E). *P <0.05, ***P < 0.001, ****P < 0.0001. Data in (A, D) are representative of two independent experiments with three replicates per condition.

We next assessed whether SARS-CoV-2 induces *PTGS2* in a more physiologically relevant cell culture system. We cultured primary human bronchial epithelial cells (HBECs) for 28 days at an air-liquid interface, which supports pseudostratified mucociliated differentiation providing an *in vitro* model of airway epithelium (15). We infected HBECs with SARS-CoV-2 at the apical surface of the culture and then performed single-cell RNA sequencing at 1, 2, and 3 days post infection (dpi) (16). As we previously reported that ciliated cells in air-liquid interface cultures are the major target of infection (16), we looked for *PTGS2* induction in this cell type. Aggregating ciliated cells across the three time points, we found that infected ciliated cells expressed higher levels of *PTGS2* compared to uninfected bystander ciliated cells (Fig. 1E), indicating that *PTGS2* is also induced by SARS-CoV-2 in a cell-intrinsic manner in ciliated cells, a physiologically relevant target cell.

To determine the relevance of these findings *in vivo*, we utilized transgenic mice expressing human ACE2 driven by the epithelial cell keratin 18 promoter (K18-hACE2) (17). As SARS-CoV-2 does not efficiently interact with mouse ACE2 (4), human ACE2-expressing mice are required to support SARS-CoV-2 infection (18–23). K18-hACE2 mice were initially developed as a model of SARS-CoV infection and have recently been demonstrated as a model of severe SARS-CoV-2 infection in the lung (17, 24). We found that intranasal infection of K18-hACE2 mice with SARS-CoV-2 led to significant upregulation of *Ptgs2* in the lung at multiple time points post infection (Fig. 1F), consistent with recent SARS-CoV-2-infected K18-hACE2 lung RNA-seq data (Fig. 1G) (24). Taken together, these results demonstrate that SARS-CoV-2 infection induces *PTGS2* in diverse *in vitro* and *in vivo* airway and lung systems, across multiple independent studies. These findings therefore suggest that COX-2/PGE_2_ signaling may be a relevant pathway for regulating SARS-CoV-2 infection and replication.

We next explored whether inhibition of the COX-2/PGE_2_ pathway could affect viral infection by regulating *ACE2* expression, as has been reported in studies of diabetic rats and intestinal organoids (7, 8). We utilized two NSAIDs, nonselective COX-1/COX-2 inhibitor ibuprofen and selective COX-2 inhibitor meloxicam, which are common in clinical use. We determined the maximum non-toxic doses of ibuprofen and meloxicam to use on Calu-3 and Huh7.5 cells (Fig. 2A-B) and validated their functionality on Calu-3 cells, which produce PGE_2_ at baseline (Fig. 2C). Treatment of Calu-3 or Huh7.5 cells with ibuprofen or meloxicam did not significantly affect *ACE2* expression (Fig. 2D-E). To test whether NSAID treatment affects *Ace2* expression in diverse tissues *in vivo*, we treated C57BL/6 mice with therapeutic doses of ibuprofen and meloxicam (25, 26), which did not lead to changes in *Ace2* expression in the lung, heart, kidney, or ileum (Fig. 3A-D). These data indicate that inhibition of the COX-2/PGE_2_ pathway by NSAIDs does not affect *ACE2* expression, and therefore susceptibility to infection, in multiple cell and tissue types *in vitro* or *in vivo*.

**Figure 2.**
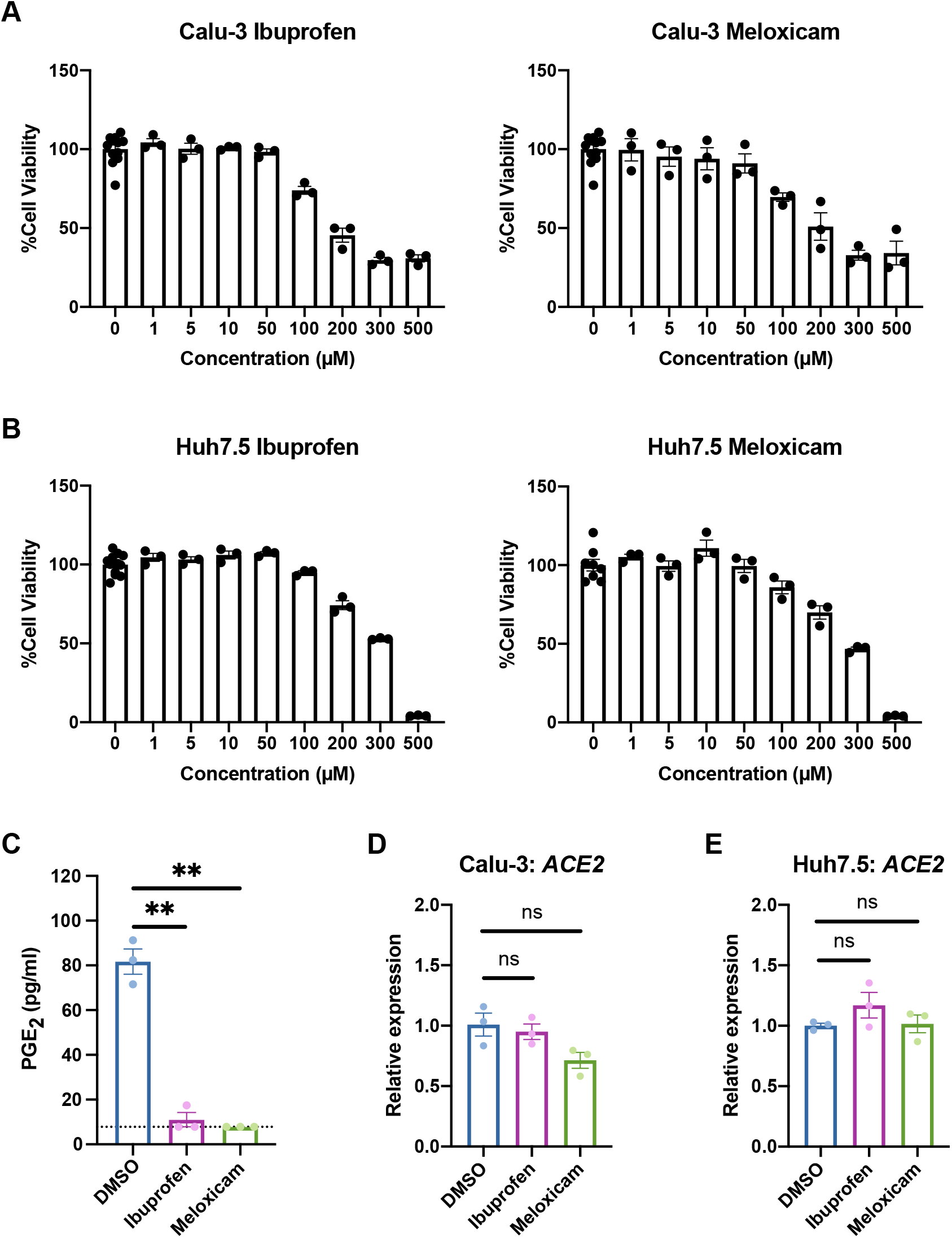
NSAID treatment does not affect *ACE2* expression in human cell lines. (A-B) Calu-3 (A) and Huh7.5 (B) cells were treated with different concentrations of ibuprofen or meloxicam for 48 hours. Cell viability was measured and calculated as a percentage of no treatment. (C) Calu-3 cells were treated with DMSO, 50 μM ibuprofen, or 50 μM meloxicam for 48 hours. Levels of prostaglandin E_2_ (PGE_2_) were measured in the supernatant. Dotted line represents limit of detection. (D-E) Calu-3 (D) and Huh7.5 (E) cells were treated with DMSO, 50 μM ibuprofen, or 50 μM meloxicam for 24 hours. *ACE2* expression was measured and normalized to *ACTB*. All data points in this figure are presented as mean ± SEM. Data were analyzed by Welch’s two-tailed, unpaired *t*-test (C-E). **P < 0.01. ns, not significant. Data in (A-E) are representative of two independent experiments with three replicates per condition.

**Figure 3.**
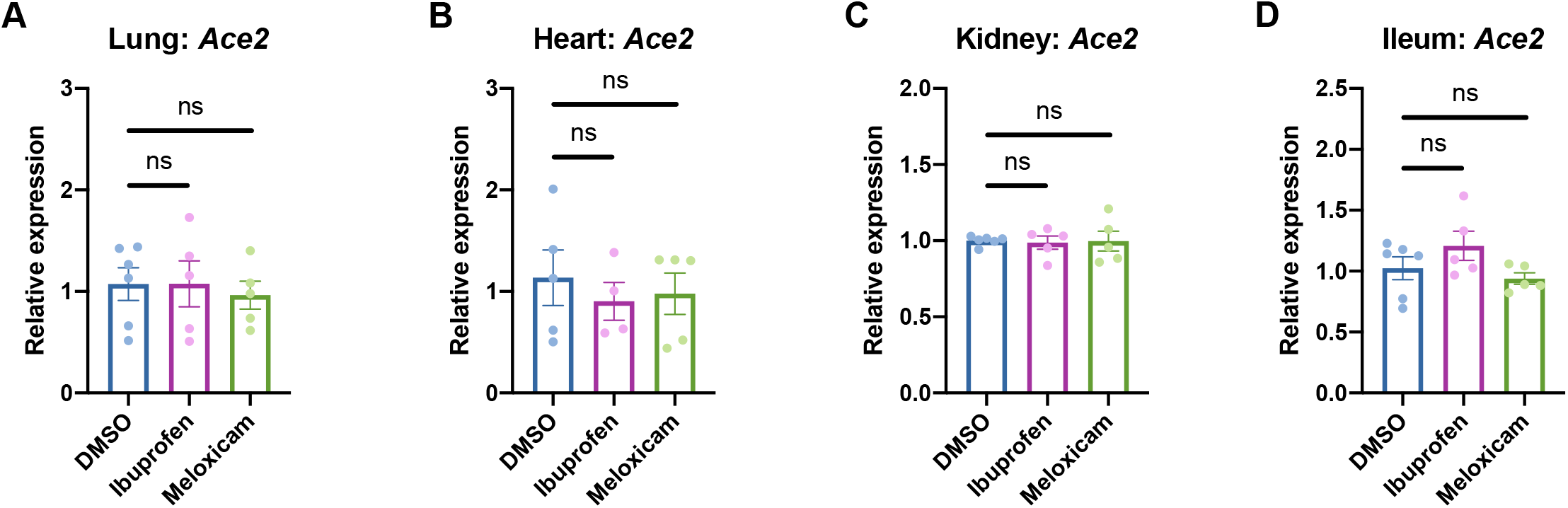
NSAID treatment does not affect *Ace2* expression in mouse tissues. (A-B) C57BL/6 mice were treated intraperitoneally with DMSO, 30 mg/kg ibuprofen, or 1 mg/kg meloxicam daily for 4 days. *Ace2* expression was measured in the lung (A), heart (B), kidney (C), and ileum (D), normalized to *Actb*. All data points in this figure are presented as mean ± SEM. Data were analyzed by Welch’s two-tailed, unpaired *t*-test (A-D). ns, not significant. Data in (A-D) are pooled from two independent experiments with four to six mice per condition.

To functionally confirm that NSAID treatment does not affect SARS-CoV-2 entry, we used a SARS-CoV-2 spike protein-pseudotyped vesicular stomatitis virus (VSV) core expressing Renilla luciferase (SARS2-VSVpp) and VSV glycoprotein-typed virus (G-VSVpp) as a control (27). Quantification of luciferase activity showed that pre-treatment of Calu-3 or Huh7.5 cells with ibuprofen or meloxicam did not significantly affect SARS2-VSVpp or G-VSVpp entry (Fig. 4A-B), confirming that NSAID inhibition of the COX-2/PGE_2_ pathway does not impact susceptibility to infection.

**Figure 4.**
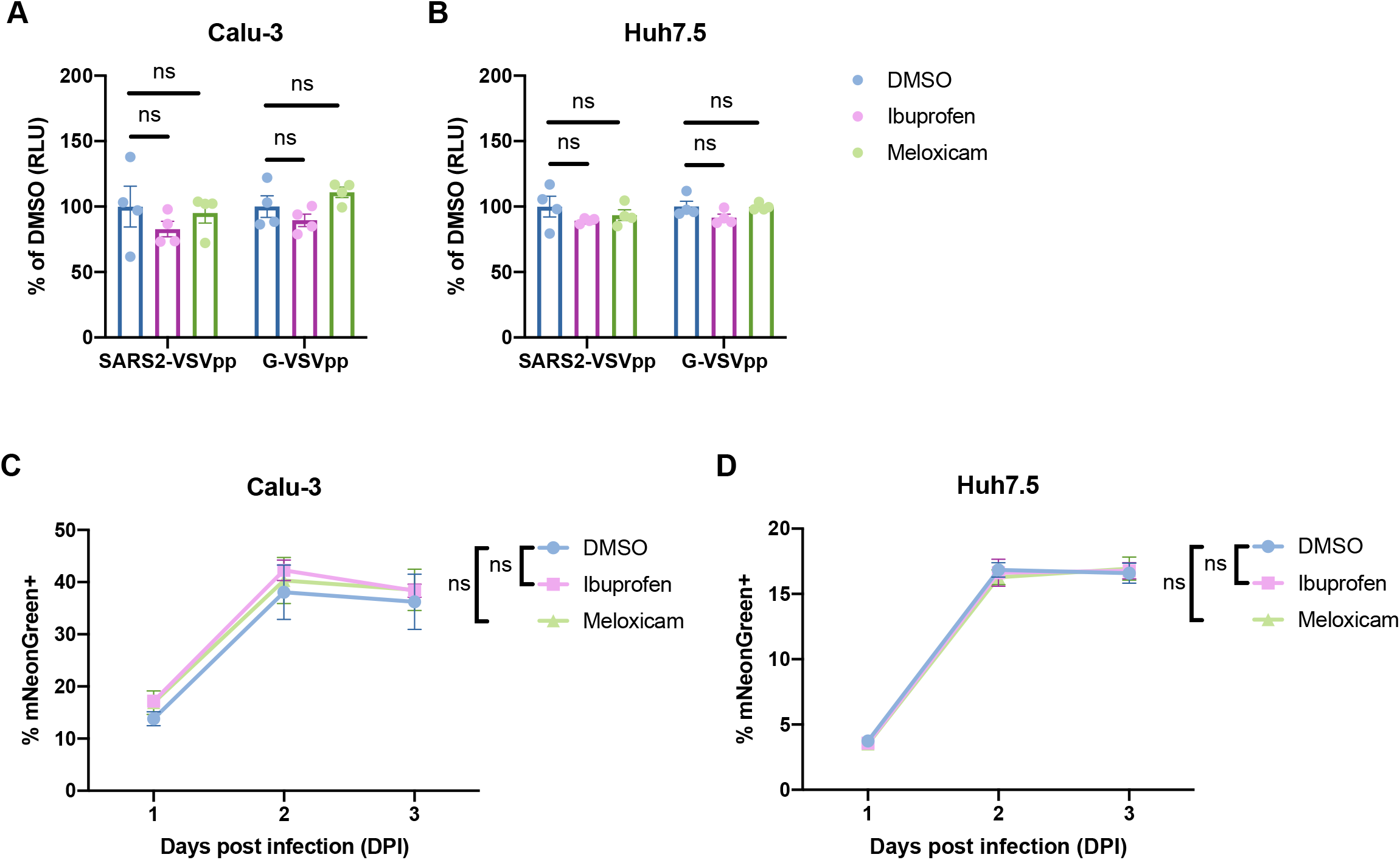
NSAID treatment does not affect SARS-CoV-2 entry or replication. (A-B) Calu-3 (A) and Huh7.5 (B) cells were pre-treated with DMSO, 50 μM ibuprofen, or 50 μM meloxicam for 24 hours and then infected with SARS2-VSVpp or G-VSVpp expressing Renilla luciferase. Luminescence was measured at 24 hours post infection (hpi) and normalized to DMSO for each infection. (C-D) Calu-3 (C) and Huh7.5 (D) cells were pre-treated with DMSO, 50 μM ibuprofen, or 50 μM meloxicam for 24 hours and then infected with mNeonGreen reporter replication-competent SARS-CoV-2 (icSARS-CoV-2-mNG) at a MOI of 1. Frequency of infected cells was measured by mNeonGreen expression at 1, 2, and 3 dpi. All data points in this figure are presented as mean ± SEM. Data were analyzed by Student’s two-tailed, unpaired *t*-test (A-B) and two-way ANOVA (C-D). ns, not significant. Data in (A-B) are representative of two independent experiments with four replicates per condition. Data in (C-D) are representative of two independent experiments with five replicates per condition.

Finally, we studied whether inhibition of the COX-2/PGE_2_ pathway affects SARS-CoV-2 replication. Viruses from several different families have been shown to induce COX-2/PGE_2_ signaling in host cells (28). The COX-2/PGE_2_ pathway is pro-viral for viruses such as porcine sapovirus, as PGE_2_ inhibits nitric oxide production, thus permitting viral replication (9). However, PGE_2_ can also be anti-viral in the case of parainfluenza 3 virus, potentially by inducing cAMP and impairing nucleic acid synthesis (29). To this end, we utilized a replication-competent SARS-CoV-2 expressing a mNeonGreen reporter (icSARS-CoV-2-mNG) to study the effect of COX-2/PGE_2_ inhibition by NSAIDs on viral replication (30). We assessed icSARS-CoV-2-mNG replication in Calu-3 cells, which upregulate *PTGS2* in response to SARS-CoV-2 infection (Fig. 1A-B), and Huh7.5 cells, which do not (Fig. 1D). Huh7.5 cells served as a control for assessing potential COX-independent effects of NSAIDs on viral replication, as has been observed with indomethacin, another NSAID, during SARS-CoV infection (31). By quantifying the percentage of mNeonGreen-expressing cells, we found that treatment of Calu-3 or Huh7.5 cells with ibuprofen or meloxicam did not impact icSARS-CoV-2-mNG replication (Fig. 4C-D). These results indicate that SARS-CoV-2 induction of the COX-2/PGE_2_ pathway in Calu-3 human lung cells does not regulate viral replication. Furthermore, ibuprofen and meloxicam do not affect SARS-CoV-2 replication in Huh7.5 cells in a COX-independent manner.

## Discussion

Given the concerns about NSAID use with COVID-19, we studied whether NSAIDs and their inhibition of the COX-2/PGE_2_ pathway affects SARS-CoV-2 entry and replication. We found that SARS-CoV-2 infection leads to *PTGS2* upregulation in diverse systems, including Calu-3 and A549 lung cancer cell lines, primary HBEC air-liquid interface cultures, and the lungs of human ACE2-expressing mice. However, inhibition of COX-2/PGE_2_ signaling with the commonly used NSAIDs ibuprofen and meloxicam did not affect *ACE2* expression in multiple cell and tissue types *in vitro* or *in vivo*, nor SARS-CoV-2 entry or replication. Our findings therefore rule out a direct effect of NSAIDs on SARS-CoV-2 infection.

An important question arising from our findings is how SARS-CoV-2 infection induces COX-2 expression. One possibility is that the pattern recognition receptor retinoic acid inducible gene-I (RIG-I), which can recognize double-stranded RNA generated during viral genome replication and transcription (32), may drive this response. Indeed, COX-2 induction by influenza A virus is RIG-I-dependent (33), and we showed here that Huh7.5 cells, which are defective in RIG-I signaling (34), do not upregulate *PTGS2* in response to SARS-CoV-2. Alternatively, SARS-CoV-2 proteins may mediate the induction of COX-2 through their complex effects on host cells. In the case of SARS-CoV, transfection of plasmids encoding either the spike or the nucleocapsid genes is sufficient to stimulate COX-2 expression (12, 13). SARS-CoV spike protein induces COX-2 expression through both calcium-dependent PKCα/ERK/NF-κB and calcium-independent PI3K/PKC∊/JNK/CREB pathways (13), while the nucleocapsid protein directly binds to the COX-2 promoter to regulate its expression (12). Any of these potential mechanisms are consistent with our HBEC scRNA-seq results demonstrating that SARS-CoV-2 increases *PTGS2* expression in a cell-intrinsic manner.

Given our finding that COX-2 signaling does not regulate viral entry or replication, the role of COX-2 induction upon SARS-CoV-2 infection remains an area for future investigation. Rather than directly affecting viral entry or replication, COX-2 induction may regulate the severe lung inflammation and injury seen in COVID-19 patients (35, 36), though it is unclear whether COX-2 would be beneficial, neutral, or detrimental to disease. COX-2 could enhance lung injury in COVID-19, as PGE_2_ has been reported to induce IL-1β and exacerbate lung injury in bone marrow transplant mice (37). Additionally, PGE_2_ stimulates fibroblast proliferation, which could underlie the fibroproliferative response to acute lung injury that results in long-lasting respiratory dysfunction (38). At the same time, PGE_2_ is critical for maintaining endothelial barrier integrity, which is disrupted in acute lung injury (39), and COX-2 has also been found to promote resolution of acute lung injury by enhancing lipoxin signaling (40). NSAID inhibition of COX-2 could therefore have complex effects on the host response to SARS-CoV-2. However, it is reassuring that retrospective studies thus far have not observed worse clinical outcomes in COVID-19 patients taking NSAIDs (41–43).

In summary, we demonstrated that SARS-CoV-2 infection induces COX-2 expression in diverse systems *in vitro* and *in vivo*. However, inhibition of COX-2 by NSAIDs did not affect viral entry or replication, suggesting NSAIDs should not be contraindicated in COVID-19 patients.

## Materials and Methods

### Cell lines

Calu-3 and Huh7.5 were from ATCC. Calu-3 cells were cultured in Eagle’s Minimum Essential Medium (EMEM) with 10% heat-inactivated fetal bovine serum (FBS), 1% GlutaMAX (Gibco), and 1% Penicillin/Streptomycin. Huh7.5 cells were cultured in Dulbecco’s Modified Eagle Medium (DMEM) with 10% heat-inactivated FBS and 1% Penicillin/Streptomycin. All cell lines tested negative for *Mycoplasma* spp.

### Generation of SARS-CoV-2 stocks

As previously described (44), SARS-CoV-2 P1 stock was generated by inoculating Huh7.5 cells with SARS-CoV-2 isolate USA-WA1/2020 (BEI Resources, NR-52281) at a MOI of 0.01 for three days. The P1 stock was then used to inoculate Vero-E6 cells, and after three days, the supernatant was harvested and clarified by centrifugation (450 × *g* for 5 min), filtered through a 0.45-micron filter, and stored in aliquots at −80°C. Virus titer was determined by plaque assay using Vero-E6 cells (44).

To generate icSARS-CoV-2-mNG stocks (30), lyophilized icSARS-CoV-2-mNG was reconstituted in 0.5 ml of deionized water. 50 μl of virus was diluted in 5 ml media and then added to 10^7^ Vero-E6 cells. Three days later, the supernatant was harvested and clarified by centrifugation (450 × *g* for 5 min), filtered through a 0.45-micron filter, and stored in aliquots at −80°C.

All work with SARS-CoV-2 or icSARS-CoV-2-mNG was performed in a biosafety level 3 facility with approval from the office of Environmental Health and Safety and the Institutional Animal Care and Use Committee at Yale University.

### Preparation of NSAIDs

Ibuprofen (I4883) and meloxicam (M3935) were purchased from Sigma-Aldrich. For cell culture experiments, ibuprofen and meloxicam were solubilized in DMSO at a stock concentration of 10 mM and then diluted in media to make working solutions. For mouse experiments, stock solutions of ibuprofen (300 mg/ml) and meloxicam (10 mg/ml) were prepared in DMSO and then diluted 100-fold in PBS to make working solutions. To determine the maximum non-toxic dose of NSAIDs to use for cell culture experiments, cells were treated with different concentrations of ibuprofen or meloxicam for 48 hours, and cell viability was measured by CellTiter-Glo® Luminescent Cell Viability Assay (Promega) following manufacturer’s instructions.

### Mice

C57BL/6J and K18-hACE2 [B6.Cg-Tg(K18-ACE2)2Prlmn/J (17)] were purchased from Jackson Laboratory. K18-hACE2 mice were anesthetized using 30% vol/vol isoflurane diluted in propylene glycol (30% isoflurane) and administered 1.2 × 10^6^ PFU of SARS-CoV-2 intranasally. C57BL/6J mice were anesthetized using 30% isoflurane and administered 30 mg/kg ibuprofen, 1 mg/kg meloxicam, or an equivalent amount of DMSO intraperitoneally in a volume of 10 ml/kg daily for 4 days. Animal use and care was approved in agreement with the Yale Animal Resource Center and Institutional Animal Care and Use Committee (#2018-20198) according to the standards set by the Animal Welfare Act. Only male mice were used due to availability.

### Analysis of RNA-seq data

We utilized RNA-seq data from recent published studies to assess the impact of SARS-CoV-2 infection on *PTGS2* expression. From GSE147507 (14), we re-analyzed the raw count data from Calu-3 and A549-ACE2 cells, comparing SARS-CoV-2 infection to matched mock controls. We performed differential expression analysis using the Wald test from DESeq2 (45), using a Benjamini-Hochberg adjusted p < 0.05 as the cutoff for statistical significance. For visualization of *PTGS2* expression, the DESeq2-normalized counts were exported and plotted in GraphPad Prism. Statistical significance was assessed using a Student’s two-tailed, unpaired *t*-test.

For analysis of HBEC air-liquid interface cultures infected with SARS-CoV-2, we utilized a previously generated catalog of differentially expressed genes that our group recently described in a preprint study (16). The differential expression table is publicly available at https://github.com/vandijklab/HBEC_SARS-CoV-2_scRNA-seq). Here, we specifically investigated *PTGS2* expression in ciliated cells, comparing infected cells to bystander cells (cells aggregated across 1, 2, and 3 dpi time points). The cutoff for statistical significance was set at adjusted p < 0.05, and the results were visualized as a volcano plot in R.

From GSE154104 (24), we re-analyzed the raw count data from the lungs of K18-hACE2 mice infected with SARS-CoV-2, performing pairwise comparisons of mice at 2 dpi, 4 dpi, and 7 dpi to 0 dpi controls (prior to infection). For visualization of *Ptgs2* expression, the DESeq2-normalized counts were exported and plotted in GraphPad Prism. Statistical significance was assessed using a Student’s two-tailed, unpaired *t*-test.

### PGE_2_ ELISA

Levels of PGE_2_ in cell culture supernatants were measured using the Prostaglandin E_2_ ELISA Kit (Cayman Chemical) following manufacturer’s instructions. Absorbance was measured at 410 nm on a microplate reader (Molecular Devices), and PGE_2_ concentrations were calculated using a standard curve.

### Quantitative PCR

Cells or tissues were lysed in TRIzol (Life Technologies), and total RNA was extracted using the Direct-zol RNA Miniprep Plus kit (Zymo Research) following manufacturer’s instructions. cDNA synthesis was performed using random hexamers and ImProm-II™ Reverse Transcriptase (Promega). qPCR was performed with Power SYBR® Green (Thermo Fisher) and run on the QuantStudio3 (Applied Biosystems). Target mRNA levels were normalized to those of *ACTB* or *Actb*. qPCR primer sequences are as follows: *ACTB* (human): GAGCACAGAGCCTCGCCTTT (forward) and ATCATCATCCATGGTGAGCTGG (reverse); *PTGS2* (human): AGAAAACTGCTCAACACCGGAA (forward) and GCACTGTGTTTGGAGTGGGT (reverse); *ACE2* (human): GGGATCAGAGATCGGAAGAAGAAAA (forward) and AAGGAGGTCTGAACATCATCAGTG (reverse); *Actb* (mouse): ACTGTCGAGTCGCGTCCA (forward) and ATCCATGGCGAACTGGTGG (reverse); *Ptgs2* (mouse): CTCCCATGGGTGTGAAGGGAAA (forward) and TGGGGGTCAGGGATGAACTC (reverse); *Ace2* (mouse): ACCTTCGCAGAGATCAAGCC (forward) and CCAGTGGGGCTGATGTAGGA (reverse).

### Pseudovirus production

VSV-based pseudotyped viruses were produced as previously described (27, 44). Vector pCAGGS containing the SARS-Related Coronavirus 2, Wuhan-Hu-1 Spike Glycoprotein Gene, NR-52310, was produced under HHSN272201400008C and obtained through BEI Resources, NIAID, NIH. 293T cells were transfected with the pCAGGS vector expressing the SARS-CoV-2 spike glycoprotein and then incubated with replication-deficient VSV expressing Renilla luciferase for 1 hour at 37°C (27). The virus inoculum was then removed and cells were washed with PBS before adding media with anti-VSV-G clone I4 to neutralize residual inoculum. No antibody was added to cells expressing VSV-G. Supernatant containing pseudoviruses was collected 24 hours post inoculation, clarified by centrifugation, and stored in aliquots at −80°C.

### Pseudovirus entry assay

3 × 10^4^ Calu-3 or 1 × 10^4^ Huh7.5 cells were plated in 100 μl volume in each well of a black-walled, clear-bottom 96-well plate. The following day, the media was replaced with 50 μM ibuprofen, 50 μM ibuprofen, or an equivalent amount of DMSO. One day later, 10 μl SARS-CoV-2 spike protein-pseudotyped or VSV glycoprotein-typed virus was added. Cells were lysed at 24 hpi and luciferase activity was measured using Renilla Luciferase Assay System (Promega) following manufacturer’s instructions. Luminescence was measured on a microplate reader (BioTek Synergy).

### icSARS-CoV-2-mNG assay

6.5 × 10^3^ Calu-3 or 2.5 × 10^3^ Huh7.5 cells were plated in 20 μl phenol red-free media containing 50 μM ibuprofen, 50 μM ibuprofen, or an equivalent amount of DMSO in each well of a black-walled, clear-bottom 384-well plate. The following day, icSARS-CoV-2-mNG was added at a MOI of 1 in 5 μl volume. Frequency of infected cells was measured by mNeonGreen expression at 1, 2, and 3 dpi by high content imaging (BioTek Cytation 5) configured with brightfield and GFP cubes. Total cell numbers were quantified by Gen5 software for brightfield images. Object analysis was used to determine the number of mNeonGreen-positive cells. The percentage of infection was calculated as the ratio of the number of mNeonGreen-positive cells to the total number of cells in brightfield.

### Statistical analysis

Data analysis was performed using GraphPad Prism 8 unless otherwise indicated. Data were analyzed using Welch’s two-tailed, unpaired *t*-test; Student’s two-tailed, unpaired *t*-test; or two-way ANOVA, as indicated. P < 0.05 was considered statistically significant.

## Data availability

All data are available as described above.

## Acknowledgements

We would like to acknowledge Benhur Lee, Pei-Yong Shi, the World Reference Center for Emerging Viruses and Arboviruses (WRCEVA), Paulina Pawlica, Joan Steitz, Michael Diamond, and BEI Resources for providing critical reagents. We thank all members of the Wilen and Eisenbarth labs for helpful discussion. We thank Yale Environmental Health and Safety for providing necessary training and support for SARS-CoV-2 research.

## Author contributions

**Jennifer S. Chen:** Conceptualization, Formal analysis, Investigation, Validation, Visualization, Writing – original draft, **Mia Madel Alfajaro:** Methodology, Investigation, **Jin Wei:** Methodology, Investigation, **Ryan D. Chow:** Formal analysis, Visualization, **Renata B. Filler:** Investigation, **Stephanie C. Eisenbarth:** Supervision, **Craig B. Wilen:** Conceptualization, Formal analysis, Funding acquisition, Resources, Supervision, Writing – original draft. All authors reviewed and edited the manuscript.

